# The complete structure of the small subunit processome

**DOI:** 10.1101/175547

**Authors:** Jonas Barandun, Malik Chaker-Margot, Mirjam Hunziker, Kelly R. Molloy, Brian T. Chait, Sebastian Klinge

## Abstract

The small subunit processome represents the earliest stable precursor of the eukaryotic small ribosomal subunit. Here we present the cryo-EM structure of the *Saccharomyces cerevisiae* small subunit processome at an overall resolution of 3.8 Å, which provides an essentially complete atomic model of this assembly. In this nucleolar superstructure, 51 ribosome assembly factors and two RNAs encapsulate the 18S rRNA precursor and 15 ribosomal proteins in a state that precedes pre-rRNA cleavage at site A1. Extended flexible proteins are employed to connect distant sites in this particle. Molecular mimicry, steric hindrance as well as protein-and RNA-mediated RNA remodeling are used in a concerted fashion to prevent the premature formation of the central pseudoknot and its surrounding elements within the small ribosomal subunit.

---

Eukaryotic ribosome assembly is a highly dynamic process involving in excess of 200 non-ribosomal proteins and RNAs. This process starts in the nucleolus where ribosomal RNAs (rRNAs) for the small ribosomal subunit (18S rRNA) and the large ribosomal subunit (25S and 5.8S rRNA) are initially transcribed as part of a large 35S pre-rRNA precursor transcript. Within the 35S pre-rRNA, the 18S rRNA is flanked by the 700-nucleotide 5’ external transcribed spacer (5’ ETS) and internal transcribed spacer 1 (ITS1), which both have to be removed during later stages of ribosome maturation^1^.

The earliest stable precursors of the small subunit (SSU), referred to here as SSU processomes, have been identified on Miller spreads by electron microscopy (EM) as terminal structures of pre-rRNA transcribed by RNA polymerase I^2,3^. In addition to the 18S rRNA precursor flanked by the 5’ ETS and parts of ITS1, these particles contain the U3 small nucleolar RNA (snoRNA) and a large number of ribosome biogenesis factors including U three proteins (Utps)^4^. A subsequent cleavage step between the 5’ ETS and the 18S (at site A1) defines the mature 5’ end of the 18S rRNA, while iterative processing at the 3’ end within ITS1 (either at site A2 or A3) separates the large and small subunit maturation pathways. A final endonucleolytic cleavage event at the mature 3’ end of the 18S rRNA (D site) results in the final 18S rRNA in the cytoplasm^5^ (Supplementary Fig. 1).

The formation of the SSU processome has previously been shown to occur in a stage-specific manner^6-8^. The 5’ ETS serves as a scaffold responsible for the recruitment of a 2-megadalton assembly called the 5’ ETS particle. Within this particle, the multi-subunit protein complexes UtpA (Utp4, Utp5, Utp8, Utp9, Utp10, Utp15 and Utp17) and UtpB (Utp1, Utp6, Utp12, Utp13, Utp18, and Utp21) are used to chaperone both the 5’ ETS and U3 snoRNA^9^. The U3 snoRNA acts as molecular guide by base-pairing with both the 5’ ETS as well as sequences within the 18S rRNA^10,11^. Together, the 5’ ETS particle components provide a platform for the ensuing steps of SSU processome formation during which the four ribosomal RNA domains of the 18S rRNA (5’, central, 3’ major and 3’ minor) are bound by a set of specific ribosome assembly factors^7,8^. Biochemical studies indicated that these factors and the 5’ ETS particle likely contribute to the independent maturation of the domains of the small ribosomal subunit^7,8^. A vital process during later stages of small subunit assembly is the formation of the central pseudoknot and its surrounding elements. Here, multiple RNA and protein elements at the interface of all four domains are juxtaposed and determine the final orientations of those domains within the mature small subunit.

Recently, the molecular architectures of SSU processomes from the thermophilic eukaryote *Chaetomium thermophilum* and the yeast *Saccharomyces cerevisiae* were determined by cryo-electron microscopy (cryo-EM) at resolutions of 7.4, 5.1 and 4.5 Ångstroms^12-14^. To obtain these related states of the SSU processome, standard growth conditions^12^, nutrient starvation^13^ as well as depletion of the exosome-associated RNA helicase Mtr4 (ref. 14) were employed. While these structures have elucidated the general architecture of this early ribosome assembly intermediate, including the positions of many ribosome assembly complexes, such as UtpA, UtpB and the U3 snoRNP, limited resolution has so far precluded the complete and unambiguous assignment of all components within the SSU processome.

Here we present the atomic structure of the *Saccharomyces cerevisiae* SSU processome at an overall resolution of 3.8 Ångstroms with local resolutions in the core near 3 Ångstroms. The combination of cryo-EM and cross-linking and mass spectrometry data has enabled us to build an essentially complete atomic model of the SSU processome containing three RNAs (5’ ETS, pre-18S and U3 snoRNA), 51 ribosome assembly factors and 15 ribosomal proteins.

## Overall structure

As previously reported, we used nutrient starvation to accumulate and purify the SSU processome^13^. This particular state may represent a storage particle or a non-productive assembly intermediate of the small ribosomal subunit in response to these stress conditions^13,15,16^. Extensive data collection and an improved 3D classification and refinement strategy enabled us to improve the resolution of the entire SSU processome to 3.8 Ångstroms (Supplementary Figs. 2, 3 and Supplementary Table 1). The core, which contains approximately 80% of the proteins present in the SSU processome, could be refined to an overall resolution of 3.6 Ångstroms. Large regions in the center of the particle are well resolved near 3 Ångstroms (Supplementary Fig. 3) and clear density can be observed for side chains and bases (Supplementary Figs. 4, 5). Focused classification and refinement resulted in significantly improved maps of more peripheral areas of the structure, such as the head region, containing the 5’ domain, or the 3’ region, containing parts of the 3’ domain (both at 4.1 Ångstroms overall resolution) (Fig. 1, Supplementary Figs. 2, 3 and Supplementary Table 1). These improved maps also feature clear side chain density for peripheral and more exposed areas (Supplementary Fig. 4) and RNA densities in different regions of the SSU processome show clear separation of nucleotides (Supplementary Fig. 5).

**Figure. 1.**
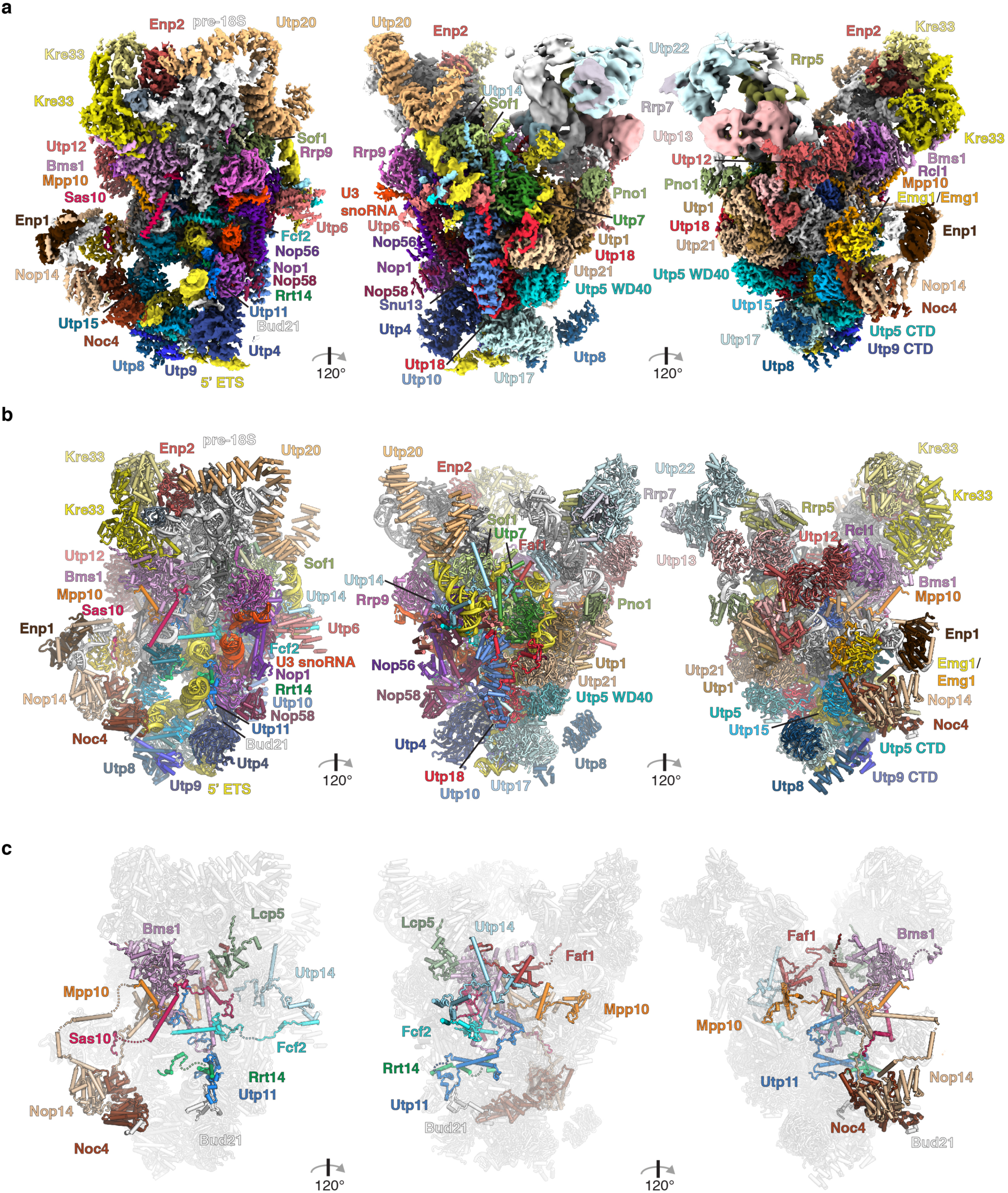
Cryo-EM reconstruction and near-complete atomic model of the *S. cerevisiae* SSU processome. **a**, Three views of a composite cryo-EM map consisting of the 3.6 Å core, the 4.1 Å head-focused, the 4.1 Å 3’ domain and the 7.2 Å central domain maps. Densities for SSU processome components are color-coded with analogous labels. Subunits of complexes are shown in shades of blue (UtpA), red (UtpB), purple (U3 snoRNP), brown (Nop14/Noc4) and light-pink (Bms1-Rcl1). Ribosomal proteins are depicted in shades of grey. RNAs are colored in yellow (5’ ETS), red (U3 snoRNA) and white (pre-18S rRNA). **b,** Cartoon representation of the atomic model of the SSU processome with orientations and colors as in (**a**). **c**, Cartoon models of newly identified extended peptides, colored as in (**a**), within the whole particle (white).

As previously shown, the central domain is largely flexible under the growth and purification conditions we used to obtain SSU processomes^13^. Extensive iterative 3D classification has enabled us to isolate one particular conformation of this domain, where density can be visualized at an overall resolution of 7.2 Ångstroms (Fig. 1a, Supplementary Figs. 2, 3 and 6).

The combined use of these maps together with cross-linking and mass spectrometry data (Supplementary Fig. 7 and Supplementary data Table 1) allowed us to unambiguously trace proteins not only in the core but also in the periphery of the particle to obtain a largely complete atomic model of the SSU processome (Fig.1, Supplementary Figs. 8-27 and Supplementary Tables 1, 2). In solvent-exposed regions, such as the central domain, where atomic resolution was not obtained, previously determined crystal structures have been fitted or poly-alanine models have been built *de novo*. We note that in these regions the sequence register of proteins is less reliable.

Like many eukaryotic superstructures, the SSU processome contains multiple β-propellers (WD40 domains)^17^ as well as helical repeat structures in addition to ribosomal components and other ribosome assembly factors (Fig. 1, Supplementary Fig. 28 and Supplementary Video 1). While helical repeat elements are frequently used to encapsulate RNA and protein elements, β-propellers perform a range of different functions. The rigid scaffold provided by seven blades of a β-propeller provides a unique platform for the individual diversification of the exposed loops, which are used for protein-protein as well as RNA-protein recognition in the 20 different β-propellers of the SSU processome. In addition, N-and C-terminal extensions provide further functional regions to interact with RNA and protein elements.

Helical repeat elements are used towards the top of the structure near the 5’ and central domains to encapsulate and stabilize RNA helices in a particular conformation. In the context of the 5’ domain, Utp20 provides a structural support for RNA expansion segments ES3A and ES3B near the Kre33 heterodimer and Enp2 (Supplementary Fig. 6). Similarly, the tetratricopeptide repeat (TPR) of Rrp5 (ref. 18), which is necessary for pre-18S processing^19,20^, is positioned in proximity to the UtpC complex^21^, Krr1 (ref.22) and the central domain (Supplementary Fig. 6). It provides a cradle to stabilize helix 24 in a different conformation with respect to the mature small subunit.

The SSU processome can be subdivided into subcomplexes (Fig. S29), which include UtpA, UtpB, UtpC, the U3 snoRNP, the Mpp10 complex, and individual proteins. UtpA and UtpB provide 16 of the twenty β-propellers within the SSU processome, with Sof1, Utp7, Enp2 and Rrp9 (ref. 23) (U3 snoRNP) providing the remaining four (Fig. 1a, b and Supplementary Fig. 28).

Limited resolution has previously prevented the correct assignment of all β-propellers within the SSU processome and in particular of Utp4 and Utp5 (UtpA) as well as Utp18 (UtpB)^12-14^. It further prohibited the tracing of intricate loops, extensions and linkers that determine the subunit-specific function of these redundant folds. By using high-resolution density maps, RNA-protein^9^ and protein-protein cross-linking data we were able to assign all subunits of UtpA and UtpB for subsequent model building (Supplementary Figs. 10-14, 30-32).

We have identified and built models of 10 previously unassigned ribosome assembly factors within the SSU processome (Fig. 1c and Supplementary Figs. 15-24). These include the 5’ ETS particle proteins Utp11, Fcf2, Sas10 and Bud21 (Fig. 2) as well as later factors such as Faf1, Lcp5, Utp14 and Rrt14 (refs 7,8). The Nop14-Noc4 sub-module^24^ was identified as a previously unassigned helical structure in the lower half of the particle. In addition, parts of Mpp10 that extend beyond the regions interacting with Imp3 and Imp4 (refs 25,26), have now been identified (Supplementary Fig. 27).

**Figure 2.**
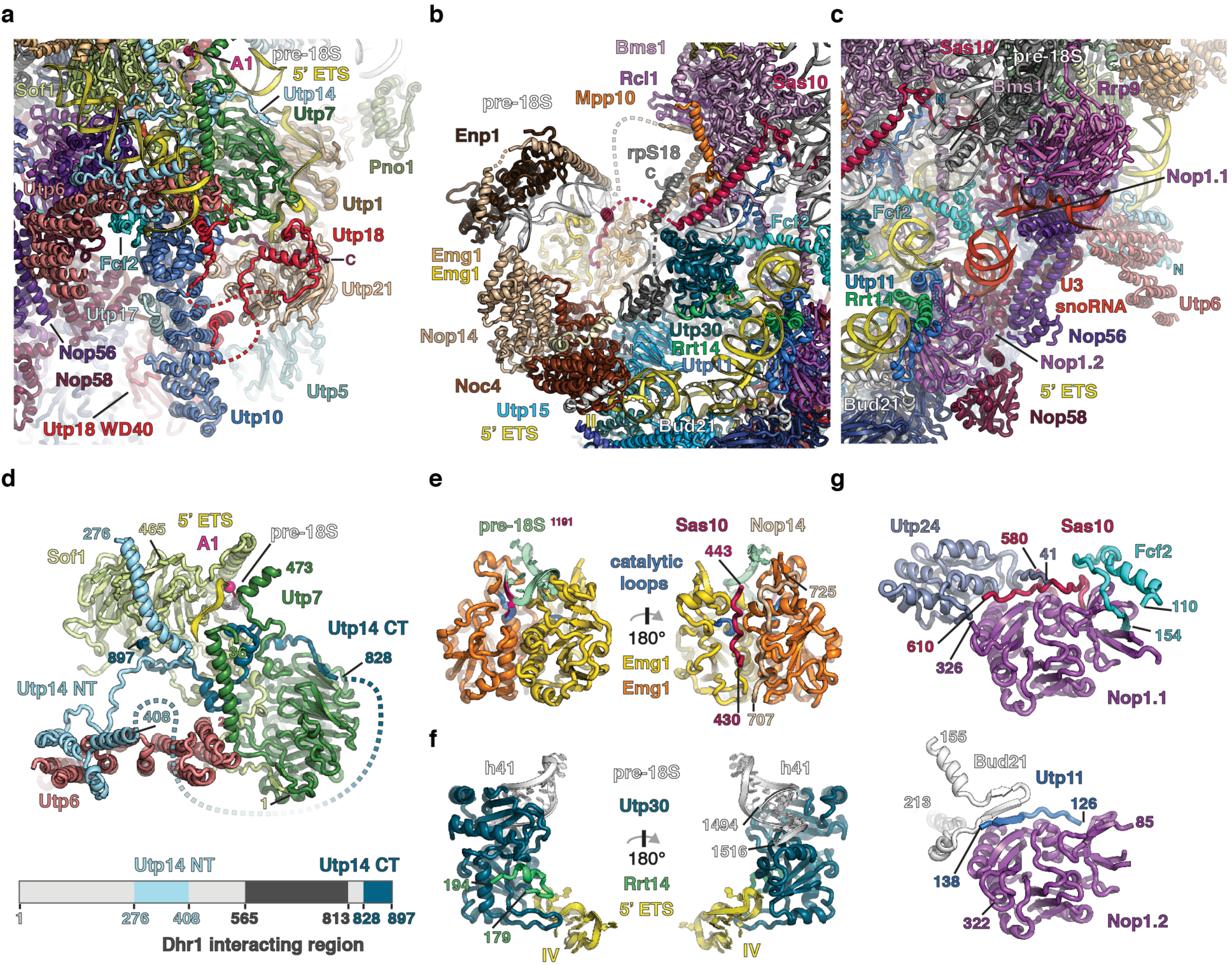
Diverse roles of peptides in the SSU processome. a-c, Views of the network of peptides traversing through the SSU processome. (**a**) Interactions of Utp18 (UtpB) with Utp10 (UtpA), Nop58 (U3 snoRNP), Utp21 and Utp6 (UtpB). Sof1, Utp7 and Utp14 organize the A1 cleavage site (pink sphere). (**b**) Nop14, Noc4 and Enp1 cover the 3’ domain of the pre-18S RNA and the methyltransferase Emg1. (**c**) Fcf2, Utp11, Bud21 and Sas10 bind to the U3 snoRNP (shades of purple) and extend through the SSU processome. **d**, Accommodation of the A1 cleavage site (pink) by Utp7, Sof1, Utp14, and Utp6. N-and C-terminal parts of Utp14 are colored light-blue and dark-blue respectively. A schematic representation of modeled parts of Utp14 (shades of blue) is shown below. **e**, Views of the Emg1 homodimer (orange, yellow) with catalytic loops (blue). Substrate pre-18S RNA (green) with target nucleotide (1191, pink) located in one active site while peptides of Sas10 (pink) and Nop14 (beige) occupy the other. **f**, Utp30 and Rrt14 recognize the pre-18S and 5’ ETS RNA. **g**, Cartoon representation of the two copies of Nop1 (Nop1.1, Nop1.2) and interacting segments of Utp24, Sas10, Fcf2, Bud21 and Utp11.

## Coordination of the 5’ ETS

UtpA forms the base of the SSU processome, where it acts as central scaffold that recognizes the first three helices (helix I-III) of the 5’ ETS. Helix I is bound by a set of loops and helical elements on top of the tandem β-propeller of Utp17 (Supplementary Fig. 31). While the β-propellers of Utp17 have functionalized top surfaces, Utp15 employs an N-terminal extension to its β-propeller and a long linker between its C-terminal domain (CTD) and WD40 domain to position helix II and to stabilize the junction between helices II and III (Supplementary Fig. 31). The WD40 domain of Utp15 and helix II further provide a binding platform for Noc4, which acts as the foundation of a lateral extension of the UtpA complex where the 3’ domain is placed (Fig 2B). This extension is additionally stabilized by Bud21, which connects Noc4 with Utp4 (UtpA), Nop1 (U3 snoRNP) and helix III that rests on top of Utp4 (Fig. 1 and Supplementary Fig. 31).

A short single stranded RNA region between helices II and III of the 5’ ETS, is coordinated by two other β-propellers located next to Utp4 and Utp17. We could unambiguously assign these WD40 domains as Utp5 (UtpA) and Utp18 (UtpB) (Fig. 1, Supplementary Figs. 12, 13).

Utp5 is integrated within UtpA through its CTD and a linker peptide, which runs along a conserved groove of Utp17. Akin to this interaction, a C-terminal peptide expansion of Utp17 contacts the β-propeller of Utp5 before binding to Utp10, the only subunit of UtpA composed solely of helical repeats (Fig. 2a). Utp5, Utp10 (UtpA), Utp21 and Utp18 (UtpB) form the junction between the UtpA and UtpB complexes. Within this junction Utp18 serves as a central nexus. The placement of the WD40 domain of Utp18 between three UtpA subunits (Utp4, Utp17 and Utp5) and near two UtpA linker regions (Utp5 and Utp15) interlocks the two largest subcomplexes of the SSU processome (Figs. 1a, b, 2a and Supplementary Figs. 31, 32).

Like Utp17, Utp18 employs extensive peptide-like motifs to facilitate protein-protein interactions (Fig. 2a). Three regions within the 230-residue N-terminus of Utp18 interact with the UtpB subunits Utp6, Utp21, the U3 snoRNP component Nop58 and Utp10 (UtpA) (Fig. 2a, Supplementary Figs. 31, 32). The first segment (residues 13-28) is employed to interact with both Utp6 and Utp10, while the second (residues 123-183) forms an intricate interface with the surface of the first β-propeller of Utp21 and a conserved C-terminal peptide of Nop58 (Fig. 2a, Supplementary Fig. 32c). Additionally, it features an Mtr4 arch interacting motif (AIM)27 which is located in a disordered region between the first and second N-terminal segment of Utp18 (Fig. 2a, Supplementary Fig. 32c). The WD40 domain of Utp18 interacts with Snu13 (U3 snoRNP) and stabilizes a single-stranded region of the 5’ ETS immediately upstream of the 3’ hinge (nucleotides 275-280). Downstream of the 3’ hinge duplex (nucleotides 293-332), the 5’ ETS is mostly single stranded with a short stem loop (nucleotides 299-308) that is stabilized by Utp21 and Utp1 on one side and Utp18 and Utp7/Sof1/Utp14 on the other side (Supplementary Fig. 32d, e). The following single stranded region between helices V and VI of the 5’ ETS (nucleotides 393-396) is bound by Imp3, which interacts with an N-terminal helix (residues 2-34) of Imp4. Together with the UtpB tetramer, Imp3 and Imp4 provide a binding surface for Mpp10.

## Roles of extended proteins in the SSU processome

Many of the newly identified proteins share striking structural properties as they all contain long linkers that are used to weave through the structure and connect distant parts within the SSU processome (Figs. 1c, 2). At the base of the structure, Bud21 acts as a connector between Utp4 and Noc4 (Supplementary Figs. 14-16) while a conserved C-terminal proline/glycine rich sequence of Rrt14 (Supplementary Fig. 19) stabilizes the interaction of the L1-domain containing (Supplementary Fig. 33) protein Utp30 with helix IV of the 5’ ETS (Figs. 1c, 2b, f). In addition, Rrt14 contacts Utp11 located on the other side of helix IV where Utp11 interacts with Nop1 and Bud21 and connects this region with the core of the SSU processome (Fig. 2c). Sas10, Utp11 and Fcf2 (Supplementary Figs. 18, 20, 21) each span at least 100 Ångstroms and are used to interconnect the U3 snoRNP with other important regions of the SSU processome (Fig. 1c, 2a-c). Strikingly, Nop1 (fibrillarin) is used as a binding platform for 5 proteins (Fcf2, Sas10, Utp24, Utp11 and Bud21) (Fig. 2g). The surfaces of the two Nop1 subunits - one located at the lower and the other at the upper end of U3 snoRNA -are used distinctively by these proteins. Here peptide backbone elements are used to form shared secondary structure elements within a β-barrel (Fcf2) or an extended β-sheet within Nop1 (Utp11, Bud21). Peptides from Sas10 and Utp24 interact similarly with Nop1 (Fig. 2g).

Utp14, Faf1, Lcp5 and Mpp10 are located near the top of the core of the SSU processome (Figs. 1c, 2a, b, d). Here, Utp14 forms a highly unusual split structure in a functionally important region near the Utp7/Sof1 dimer and the A1 cleavage site (Fig. 2a, d and Supplementary Fig. 23). An N-terminal segment of Utp14 is used to connect Sof1 with Utp6 while a separate C-terminal segment of Utp14 links Utp7 with Sof1. Several hundred residues of Utp14 connect these two fragments. Close to the C-terminal segment, this sequence contains the binding site for Dhr1, the essential DEAH box helicase that is responsible for displacing U3 snoRNA from early ribosome assembly intermediates^28,29^. Faf1 is positioned in proximity to Utp14 and Utp7 and directly interacts with U3 snoRNA and pre-18S rRNA near the A1 cleavage site (Fig. 1b, c, Supplementary Fig. 22). Lcp5 interacts extensively with rRNA of the 5’ domain and rpS9 (Supplementary Fig. 24) whereas Mpp10 extends from Bms1 via Imp4 and Imp3 to the UtpB complex and helix 44. Several of the newly identified proteins (Fcf2, Sas10, Mpp10, Utp11 and Nop14) interact with the centrally positioned GTPase Bms1 (ref. 30) (Fig. 1 and Supplementary Figs. 26, 34). In addition to the translational GTPase fold, Bms1 contains a long C-terminal helix, which is used to anchor this enzyme in the core of the SSU processome.

In contrast to most of the other newly identified proteins, which have no visible globular domains, Nop14 contains a core helical repeat in addition to its long linker peptides (Supplementary Fig. 17). By directly binding to a second repeat protein, Noc4, the core domain of Nop14 is further expanded into a larger repeat structure. This composite helical repeat serves as a lateral structural extension of the UtpA complex and provides a scaffold for Enp1, Emg1 and parts of the 3’ domain of the 18S rRNA (Fig. 2b, Supplementary Figs. 16, 17), which later forms the beak structure in the mature small subunit. The repeat of Nop14 is flanked by N-and C-terminal extensions. A 75 amino acid long C-terminal helix docks Nop14 into an opening between the Mpp10-Imp4 dimer and the Bms1-Rcl1 complex^31^ and points its C-terminal end towards the central cavity between the central and the 5’ domains. The N-terminal extension of Nop14 binds Enp1, which caps the 3’ domain of the 18S rRNA. Surprisingly, the N-terminal segment of Nop14 loops around this entire region and extends back to where the C-terminal long helix is located (Supplementary Fig. 17). Similar to Bms1, terminal extensions are used to fully integrate the Nop14/Noc4 complex within the center of the SSU processome (Figs. 1c, 2b). In addition to stabilizing Enp1 on top of the rRNA, this arrangement also positions the rRNA substrate in the active site of one of the Emg1 dimers. Peptides from Nop14 and Sas10 are used to provide structural support for the dimeric Emg1 while at the same blocking the active site of one of its subunits. This enforces a structural asymmetry of the two Emg1 methyltransferase subunits so that only one active site is available for the methylation of base 1191 of the 3’ domain (Fig. 2e).

Near the base of helix 41 of the 18S rRNA, we have identified rpS18, a ribosomal protein that is already positioned in a near-mature configuration with respect to the 18S rRNA (Fig. 2b and Supplementary Fig. 25). The C-terminus, which binds elements of the beak structure in the mature 40S subunit, adopts a different conformation in the context of the SSU processome, where this structure has not yet formed. Here, the C-terminus of rpS18 is stabilized by the long C-terminal helix of Nop14 as well as domain IV of Bms1 and a linker region of Mpp10 (Fig. 2b).

## Protein-assisted RNA remodeling by U3 snoRNA

U3 snoRNA occupies a central position within the SSU processome and reaches from the outside into the core of the particle (Fig. 3a). By base pairing with its 5’ and 3’ hinges to nucleotides within the 5’ ETS, it rigidifies the structural scaffold provided by the 5’ ETS, which has been described previously13. The 5’ end of U3 snoRNA reaches further into the center of the SSU processome and base pairs with two regions of the pre-18S rRNA. A similar base pairing pattern was recently proposed for the SSU processome captured in a state upon Mtr4 depletion^14^. While Box A (U3 nucleotides 16-22) is base paired with nucleotides 9-15 of the pre-18S rRNA and organizes the pre-18S 5’ end near the A1 cleavage site, Box A’ (U3 nucleotides 3-13) is base paired with nucleotides 1111-1122 and re-organizes this region that is later in proximity to the central pseudoknot (Fig. 3a, Supplementary Fig. 35). A range of ribosome assembly factors is responsible for the stabilization of the four RNA duplexes that U3 snoRNA forms with the 5’ ETS and the 18S precursor. Towards the outside of the particle, three proteins of the UtpB complex (Utp18, Utp21 and Utp1) are stabilizing the 3’ hinge. While Utp18 is involved in rigidifying the junctions between helices II, III and IV of the 5’ ETS, Utp21 and Utp10 (UtpA) form a clamp around the 3’ hinge (Supplementary Fig. 32). Utp1 is used to bind the single stranded regions of both U3 snoRNA and 5’ ETS with two long structured loops (residues 556-580 and 616-680). These loops act as a rudder, thereby separating the 5’ ETS and the U3 snoRNA (Supplementary Fig. 32e).

**Figure 3.**
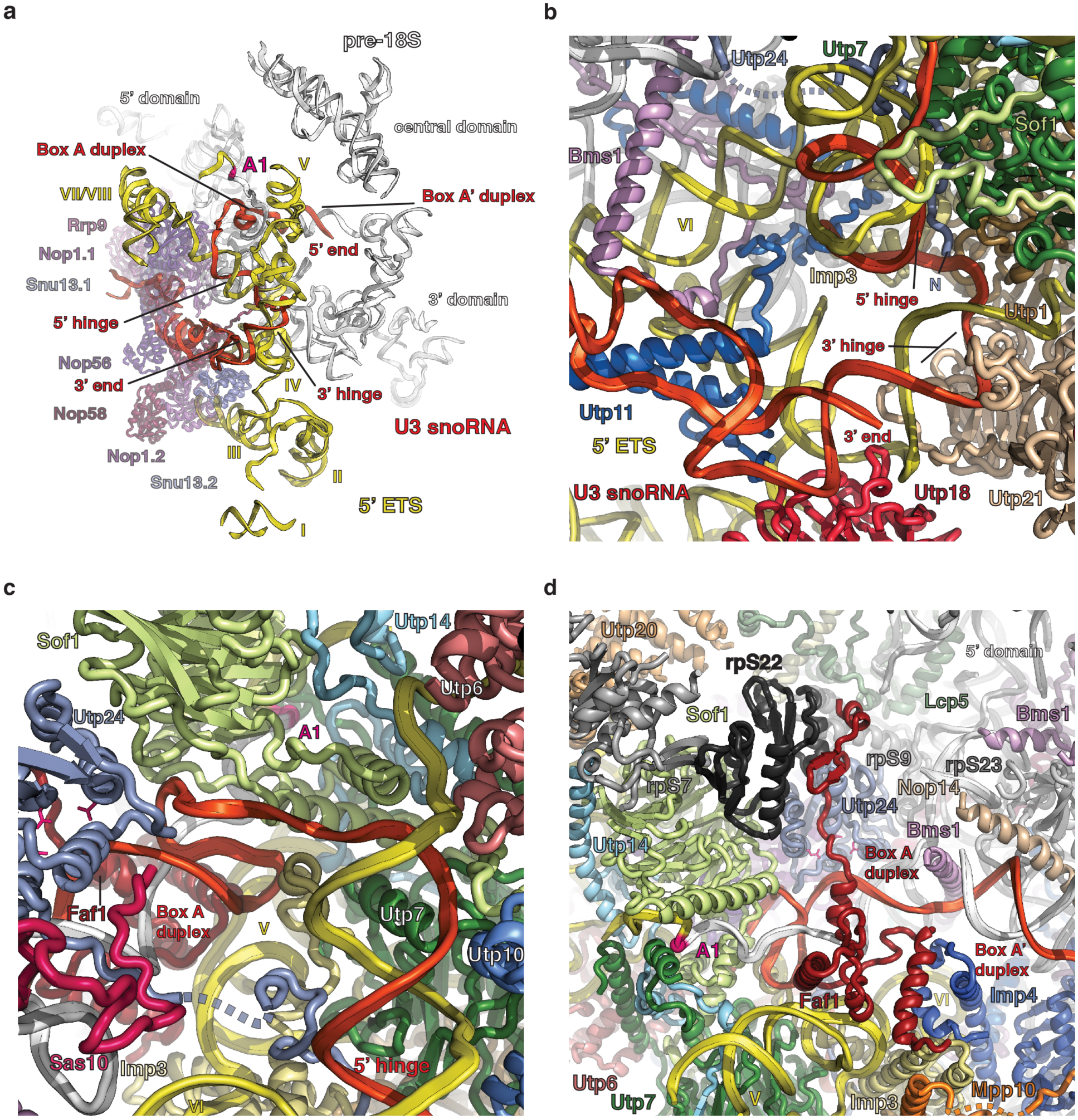
U3 snoRNA-mediated RNA remodeling. **a**, Overview of the U3 snoRNP proteins (purple), U3 snoRNA (red) and their interactions with the 5’ ETS (yellow) and pre-18S RNA (white). **b**, View of protein elements stabilizing the 3’ and 5’ hinges. **c**, View of proteins organizing the 5’ hinge and the Box A duplex. **d**, Overview of the Box A and Box A’ duplexes of U3 snoRNA with pre-18S RNA. (**a**-**d**) Proteins and RNAs are color-coded. The N-terminus of Utp24 is indicated with an N. Helices of the 5’ ETS are labeled with roman numbers. Important functional sequence elements of the U3 snoRNA (5’ hinge, 3’ hinge, Box A duplex, Box A’ duplex) and visible domains of the 18S rRNA (3’, 5’ and central domain) are indicated. The cleavage site A1 is shown as pink sphere. Active site residues of Utp24 are highlighted as pink sticks.

A short loop of U3 snoRNA, and the 5’ hinge are coordinated by Imp3, Utp11, Bms1 and the N-terminus of Utp24 (Fig. 3b). Upstream of the 5’ hinge, U3 snoRNA is stabilized by the Sof1/Utp7/Utp14 complex, which also provides a composite binding site for a single stranded RNA that contains the A1 cleavage site (Fig. 2d, 3c, d). The Box A and Box A’ duplexes are organized by the long C-terminal helices of Nop14 and Bms1 as well as Utp24, Faf1 and rpS23. Importantly, the captured state of the SSU processome contains an A0-cut precursor^13^ in a pre-A1 cleavage state where Utp24 is positioned close to its substrate but cleavage has not yet occurred. Faf1 is positioned between helix V of the 5’ ETS and the Box A duplex near Imp3 and Imp4. A linker of Faf1 locks the Box A duplex in place and interacts with rpS22 on the opposite side, thereby occluding access to the active site of the nuclease Utp24 (Fig. 3d).

## RNA remodeling prevents central pseudoknot formation

In the context of the SSU processome, Lcp5 and Sas10 are multi-functional proteins. In addition to Utp18, which contains a peptide that can interact with the exosome-associated helicase Mtr4 (ref. 27), Lcp5 and Sas10 contain exosome interaction motifs (Sas10 domains)^32^.

Lcp5 is positioned next to rpS9 and the 5’ domain, a location, which is occupied by expansion segment ES6D of the central domain in the mature small subunit (Fig. 4). In particular, the presence of expansion segment ES6D near rpS9 indicates a mature conformation of the 5’ and central domains with respect to each other, which is not the case in the context of the SSU processome.

**Figure 4.**
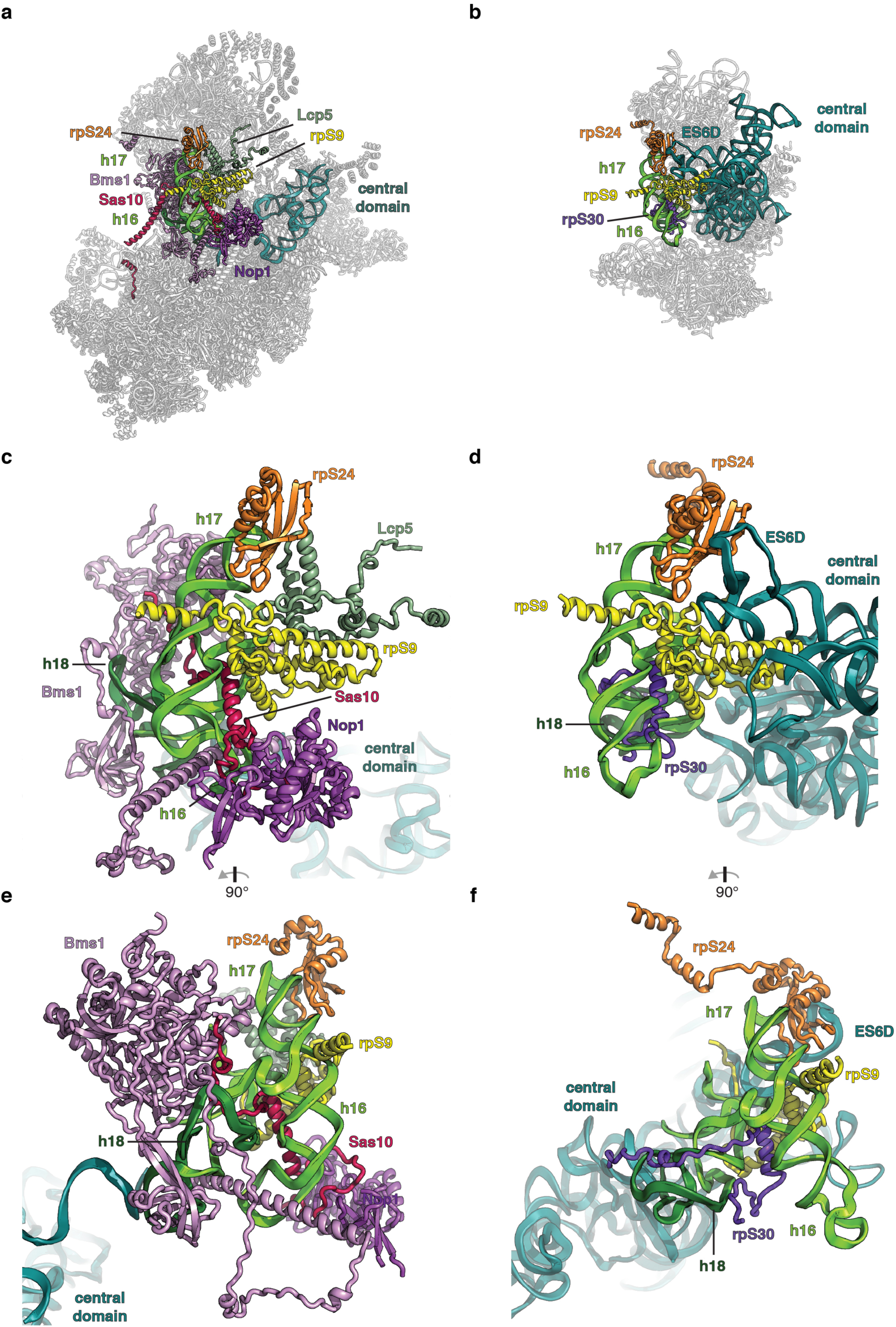
Steric hindrance and molecular mimicry prevent premature ribosomal-RNA folding. **a**, Conformations of helices 16, 17 (green) and 18 (dark green) of the 5’ domain, and the central domain (teal) of the pre-18S RNA within the SSU processome (grey). Interacting ribosome assembly factors and ribosomal proteins are shown and color-coded. **b**, Conformation of the same elements as in (**a**) in the context of the mature small ribosomal subunit (PDB 4V88) (ref. 33). **c**,**d**, Detailed views of the conformations of helix 16 and the central domain in the SSU processome (**c**) and the small ribosomal subunit (**d**). Sas10 mimics rpS30 and occupies its binding site on helix 16. Lcp5 blocks the central domain from occupying its mature position by steric hindrance while Bms1 bends helix 16. **e,f**, Orthogonal views to panel (**c**,**d**).

In contrast to Lcp5, the ordered parts of Sas10 are more elongated. Sas10 contains a solvent exposed N-terminal Sas10 domain, blocks the active site of Emg1 with a short peptide and bridges between the 5’ and 3’ domains with a long helix near Utp30 (Fig. 1b, c). In addition, it interacts with the U3 snoRNP through its C-terminus and is involved in RNA/protein remodeling by adopting a similar conformation, and occupying the same RNA binding site as rpS30 in the mature small subunit, close to rpS9 (Figs. 1b, c, 2b, c, and 4).

The continued presence of Sas10 or Lcp5 is therefore mutually exclusive with the mature small subunit conformation in terms of protein and RNA occupancy respectively and may therefore signal an incomplete or faulty assembly state during later stages of ribosome assembly.

Within the SSU processome, the four structured domains of the 18S rRNA (5’ central, 3’ major and 3’ minor domain) are segregated into different regions of the particle, thereby facilitating their separate maturation^13^. RNA elements that are positioned in the vicinity of the central pseudoknot in the mature small subunit^33^ are distinctly separated in the SSU processome (Fig 5a, b, Supplementary Fig. 6). This is organized through multiple mechanisms. RNA mediated chaperoning of sequences close to the central pseudoknot in the 5’ and central domains is accomplished through base pairing with U3 snoRNA boxes A and A’ (Fig. 3a, c, d). As a consequence of these interactions, sequences vicinal to the central pseudoknot, such as helix 27, adopt different conformations in the SSU processome, where a new stem loop forms between Rcl1 and Utp12 (Fig. 5a).

**Figure 5.**
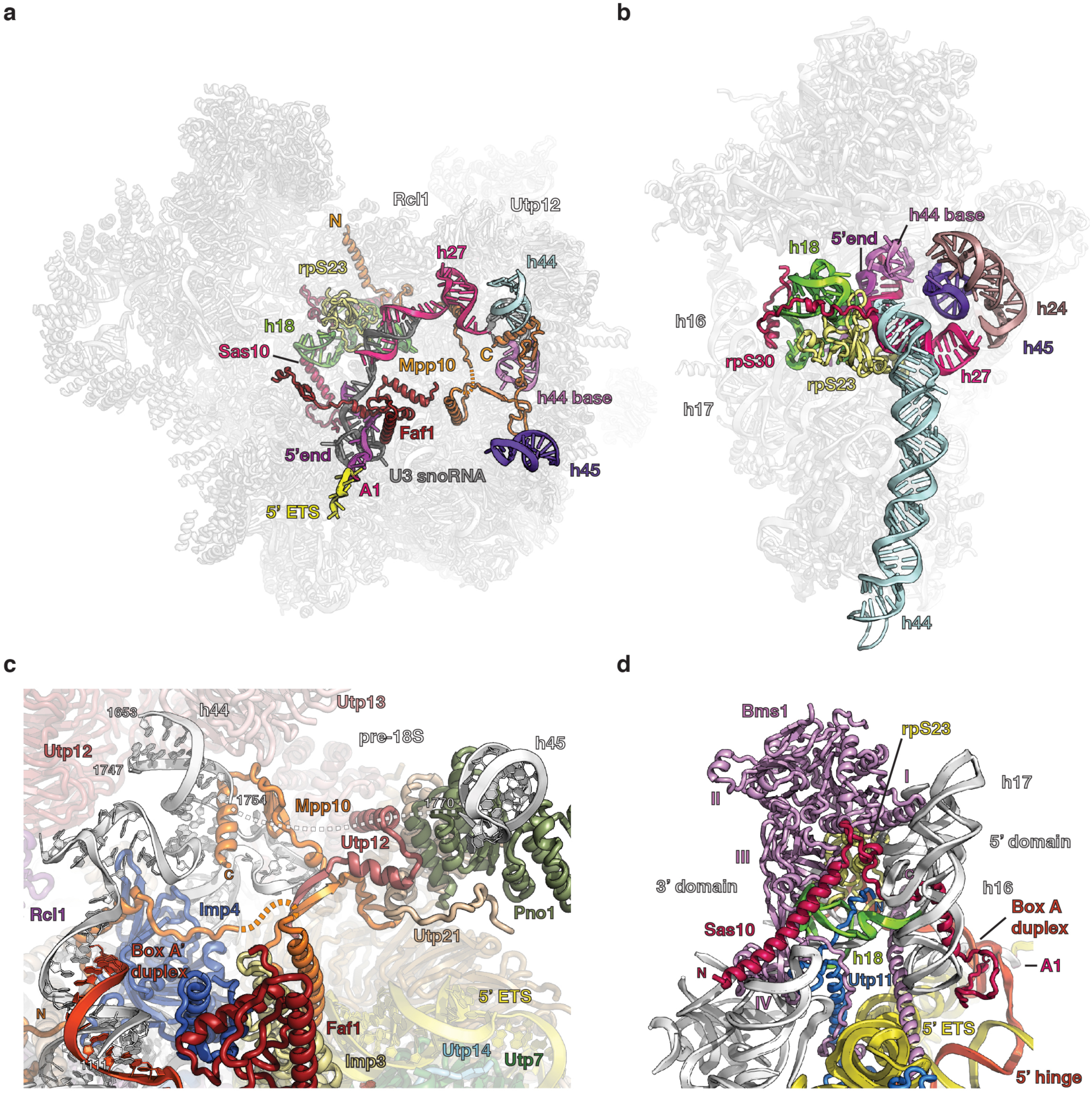
RNA remodeling prevents central pseudoknot formation. **a**, The central pseudoknot and 18S rRNA elements in its vicinity are shown in color in their immature positions in the SSU processome (grey) and labeled with their corresponding mature 18S rRNA helix (h) number. Chaperoning RNA, ribosomal proteins and ribosome assembly factors are color-coded and the A1 cleavage site is highlighted in pink. Termini of Mpp10 are indicated with N and C respectively. **b**, Color-coded RNA elements close to the central pseudoknot in the mature small ribosomal subunit (grey) labeled as in (**a**) (PDB 4V88) (ref. 33). **c**, Mpp10 (orange) and its interactions with pre-18S RNA (white). Elements of the 18S rRNA (helices 44 and 45) and U3 snoRNA (Box A’ duplex, red) are labeled. Nucleotide positions of the pre-18S RNA are indicated by white numbers. **d**, Bms1-mediated remodeling of helix 18 (h18, green) of the pre-18S RNA (white). Domains of Bms1 (purple) are numbered with roman letters. Structural elements (h16, h17) and domains of the 18S rRNA (3’ domain, 5’ domain) as well as the U3 snoRNA (5’ hinge, Box A duplex) are labeled. Other factors assisting in the remodeling (Utp11, Sas10) and the ribosomal protein rpS23 are shown.

Mpp10 plays a central role in the remodeling of nucleotides close to helices 44 and 45. A partial unwinding of the region upstream of helix 44 results in an RNA loop (nucleotides 1628-1639) that is stabilized by Mpp10 (Fig. 5c). Due to this partial unwinding, 16 nucleotides of the opposite strand (nucleotides 1755-1769) are available to serve as a linker to helix 45, which is positioned 60 Ångstroms away on top of Pno1 that is held in place by Utp1 and Utp21 of the UtpB complex (Fig. 5c).

A second important location for protein-mediated RNA remodeling is the binding site of ribosomal protein rpS23, which is positioned in the mature small subunit close to all other remodeled RNA elements next to helix 18 (Fig. 5b). In the SSU processome, conserved elements of Bms1, Utp11 and Sas10 are employed in a concerted fashion to dramatically remodel helix 18 (h18, nucleotides 558-590) of the 18S rRNA (Figs. 4c-f, 5d). The C-terminal linker region and domain IV of Bms1 together with the conserved N-terminal segment of Utp11 and a conserved linker region of Sas10 stabilize the remodeled RNA as well as rpS23, which is located in proximity to domains I-III of Bms1. Combined with the chaperone functions of Utp20 and Rrp5 in the 5’ and central domains (Supplementary Fig. 6), these examples highlight the precise and elaborate mechanisms that are employed to control tertiary interactions between ribosomal RNA domains.

## Emerging eukaryotic regulatory mechanisms

The evolution of eukaryotes has been accompanied by the emergence of dedicated multi-protein complexes that are uniquely suited to fulfill specific tasks. Eukaryotic ribosome biogenesis is catalyzed by a complex assembly of specialized protein factors, many of which have no counterparts in prokaryotes. The atomic structure of the SSU processome provides a molecular snapshot of approximately a quarter of these 200 factors. These proteins share an essential role in providing an additional level of control to ribosome biogenesis. This is achieved by concerted RNA remodeling that prevents the premature formation of the junction between all four 18S rRNA domains.

Defects in ribosome assembly are associated with a set of human diseases, called ribosomopathies^34,35^, of which some can now be mapped in a structural context (Supplementary Fig. 36). In particular, two residues near the Bms1-rpS23 interface have been implicated in human diseases. While one residue is positioned in a dynamic loop of rpS23 (ref. 36) (Supplementary Fig. 36d, e, g), the other is located within Bms1 (ref. 37) and interacts with rpS23 (Supplementary Fig. 36f).

A second function of the eukaryotic ribosome biogenesis machinery is the encapsulation and guided stabilization of a series of early ribosome assembly intermediates, which ultimately result in the mature small subunit^9,38^. The formation of the SSU processome may be directed by an initially flexible set of peptide interactions. As SSU processome maturation progresses, the number of these peptide interactions likely increases and results in this stable nucleolar superstructure.

The current state of the SSU processome highlights the need for extensive RNA and protein remodeling by specialized enzymes. This is exemplified by the numerous base pairing interactions between U3 snoRNA and 18S rRNA, which need to be unwound by enzymes such as Dhr1 to facilitate further maturation of the small ribosomal subunit. In addition, the order of the catalytic reactions performed by enzymes within the SSU processome, such as the acetyltransferase/helicase Kre^33^, the methyltransferase Emg1, the GTPase Bms1 and the nuclease Utp24, is still unknown.

Ultimately, the separation of the 5’ ETS, U3 snoRNA and 18S rRNA precursor represents a major hurdle. To accomplish this, 5’ ETS degradation and release of the 18S rRNA precursor may be coupled via the three proteins (Utp18, Lcp5 and Sas10) that can interact with the RNA surveillance machinery. This would provide an elegant mechanism to combine the recycling of ribosome assembly factors, release of a pre-40S particle and quality control.

## METHODS

### Purification of the small subunit processome

The small subunit processome was purified as previously described13 from a *Saccharomyces cerevisiae* BY4741 strain harboring a TEV-protease cleavable C-terminal GFP-tag on Utp1/Pwp2 (Utp1-3myc-TEV-GFP-3FLAG) and a second streptavidin binding peptide tag on Kre33 (Kre33-sbp). Yeast cultures were grown to an optical density of 0.6-1 in full synthetic media containing 2% raffinose (w/v) at 30 °C prior to the addition of 2% galactose (w/v) and subsequently grown to saturation. Cells were harvested by centrifugation at 4000 x *g* for 5 minutes at 4 °C. The pellet was washed with ice cold ddH2O, first without, then supplemented with protease inhibitors (E-64, Pepstatin, PMSF). Washed cells were flash frozen in liquid nitrogen and lysed by cryogenic grinding using a Retsch Planetary Ball Mill PM100.

The obtained yeast powder was resuspended in buffer A (50 mM Tris-HCl, pH 7.7 (20 °C), 150 mM NaCl, 1 mM EDTA, 0.1% Triton-X100, PMSF, Pepstatin, E-64), cleared by centrifugation at 4 °C, 40,000 x *g* for 10 min and incubated with anti-GFP nanobody beads (Chromotek) for 2 hours at 4 °C. Beads were washed three times in buffer A before bound protein complexes were eluted through TEV-protease cleavage (1 hour, 4 °C). The eluted supernatant was subjected to a second affinity purification step by incubation with streptavidin beads (Sigma) in buffer B (50 mM Tris-HCl, pH 7.7 (20 °C), 150 mM NaCl, 1 mM EDTA) for 1 hour at 4 °C.

For cryo-EM grid preparation, the streptavidin beads were subsequently washed four times in buffer B and the SSU processome was eluted in the same buffer, supplemented with 5 mM D-biotin. For protein-protein cross-linking analysis, the beads were washed in buffer C (50 mM HEPES-NaOH, pH 7.7 (4 °C), 150 mM NaCl, 1 mM EDTA) and eluted in buffer C supplemented with 5 mM D-biotin.

### Cryo-EM grid preparation

Grids were prepared from separate SSU processome purifications to collect four separate datasets (ds1-ds4). First, the sample (in 50 mM Tris-HCl, pH 7.7 (20 °C), 150 mM NaCl, 1 mM EDTA, 5 mM D-biotin) at an absorbance of 1.2 mAU to 2.4 mAU at 260 nm (Nanodrop 2000, Thermo Scientific) was supplemented with 0.03% Triton-X100 (ds1) or 0.1% Triton-X100 (ds2, ds3, ds4). Subsequently, 3.5 to 4 μl sample was applied onto glow-discharged grids (30 seconds at 50 mA) and flash frozen in liquid ethane using a Vitrobot Mark IV (FEI Company) (100% humidity, blot force of 0 and blot time 2 s). The grids for ds1, ds3 and ds4 were prepared using lacey-carbon grids (TED PELLA, Inc, Prod. No. 01824), while Quantifoil R 1.2/1.3 Cu 400 mesh grids (Agar Scientific) were used for ds2. Both grid types contained an ultra-thin carbon film.

### Cryo-EM data collection and image processing

10,029 micrographs were collected in four sessions (ds1 - ds4) on a Titan Krios (FEI Company) operated at 300 kV, mounted with a K2 Summit detector (Gatan, Inc.). The micrographs from ds1 have been obtained previously^13^ and have been included in this larger dataset and re-processed together with ds2, ds3 and ds4 (Supplementary Fig. 2). SerialEM^39^ was used to automatically acquire micrographs with a defocus range of 0.6 to 2.6 μm at a pixel size of 1.3 Å. Movies with 32 frames were collected at a dose of 10.5 electrons per pixel per second over an exposure time of 8 seconds resulting in a total dose of 50 electrons per Å2. Data collection parameters can be found in Supplementary Table 1.

All 32 movie frames were gain corrected, aligned and dose weighted using MotionCor2 (ref. 40). CTFFIND 4.1.5 was used for estimating the contrast transfer function (CTF)^41^. Manual inspection and the elimination of micrographs with bad CTF fits or drift reduced the number of micrographs from 10,029 to 8,406. Particles were first picked automatically using the RELION-2.0 (ref. 42) Autopick feature and then subjected to manual curation which yielded a total of 772,120 particles. These particles were extracted with a box size of 400 pixels (520 Å) for 3D classification. 2D classification was skipped to retain rare views of the particles. 3D classification was performed with 5 classes using EMD-8473 (ref. 13), low-pass filtered to 60 Å, as an input model (Supplementary Fig. 2). This 3D classification produced two good classes, both combined containing a total of 284,213 particles. Auto-refinement and post processing in RELION-2.0 yielded a map with an overall resolution of 3.8 Å with large areas in the center of the particle near 3 Å local resolution (Supplementary Fig. 3a). A focused refinement using a mask encompassing the “core” region further improved the quality of the map in the best resolved areas of the particle (Supplementary Figs. 2, 3b). By using a mask encompassing UtpA, the Nop14/Noc4 heterodimer and the 3‘ domain, we were able to obtain continuous density for the RNA and the Enp1 repeat protein and also improved the density for UtpA significantly (Supplementary Figs. 2, 3c).

To improve the peripheral areas near the top of the particle, iterative 3D classification (first without, later with image alignment) was done using a mask around the head region, including the Utp20 helical repeat protein. The best class from these classification steps was used for subsequent 3D refinement without mask. This strategy yielded a reconstruction at an overall resolution of 4.1 Å, with good density throughout the particle and an improvement in the head region (Supplementary Figs. 2, 3d).

Similarly, iterative focused classifications (with and without image alignment) were used for the central domain, where one specific conformation was isolated. This conformation is present in 15% of the particles used to generate overall map 1. Focused 3D refinement lead to an improvement of the resolution of this domain to 7.2 Å, allowing for the docking of crystal structures (Supplementary Figs. 2, 3e, 6). Local resolution was estimated using Resmap^43^. All computation was performed on a single Thinkmate SuperWorkstation 7048GR-TR equipped with four NVIDIA QUADRO P6000 video cards, 2 x Twenty-two Core Intel Xeon 2.40 GHz Processors, and 512 GB RAM.

### Model building and refinement

The poly-alanine model of the SSU processome provided by PDB 5TZS (ref. 13) served as a starting scaffold for the building of the current model. SSU processome proteins and RNA were either *de novo* modeled or, if applicable, available crystal structures were docked and manually adjusted. Phyre models were used as initial template for the model building of some proteins^44^. All model building was done in *Coot*^45^. A complete list of templates, crystal structures and maps used to build the model can be found in Supplementary Table 2.

The model was refined against overall map 1 (3.8 Å) in PHENIX with phenix.real_space_refine and secondary structure restraints for proteins and RNAs^46^. Model statistics can be found in Supplementary Table 1.

### Map and model visualization

Structure analysis and figure preparation was performed using PyMOL Molecular Graphics System, Version 1.8 Schrödinger, LLC and Chimera^47^. Molecular graphs and analyses were also performed with UCSF ChimeraX, developed by the Resource for Biocomputing, Visualization, and Informatics and the University of California, San Francisco (supported by NIGMS P41-GM103311).

### DSS cross-linking mass spectrometry sample preparation and analysis

Final elution fractions of tandem-affinity purified SSU processome samples (in 50 mM HEPES-NaOH, pH 7.7 (4 °C), 150 mM NaCl, 1 mM EDTA, 5 mM D-biotin) with an absorbance of 0.5 mAU at 260 nm (Nanodrop 2000, Thermo Scientific) were pooled (total volume 3 ml) and split into twenty 150-μl cross-linking reaction aliquots.

To each aliquot, 1.5 μl of disuccinimidylsuberate (DSS; 50 mM in DMSO, Creative Molecules Inc.) was added to yield a final DSS concentration of 0.5 mM and samples were cross-linked for 30 minutes at 25 °C with 450 rpm constant mixing. The reactions were quenched with 50 mM ammonium bicarbonate (final concentration) and precipitated by adding methanol (Alfa Aesar, LC-MS grade) to a final concentration of 90% followed by overnight incubation at -80 °C. Precipitated cross-linked SSU processomes were collected in one tube by repeated centrifugation at 21,000 x *g*, 4 °C for 30 minutes. The resulting pellet was washed three times with 1 ml cold 90% methanol, air-dried and resuspended in 50 μl of 1X NuPAGE LDS buffer (Thermo Fisher Scientific).

DSS cross-linked SSU processomes in LDS buffer were reduced with 25 mM DTT, alkylated with 100 mM 2-chloroacetamide, separated by SDS-PAGE in three lanes of a 3-8% Tris-Acetate gel (NuPAGE, Thermo Fisher Scientific), and stained with Coomassie-blue. The gel region corresponding to cross-linked complexes was sliced and digested overnight with trypsin to generate cross-linked peptides. After digestion, the peptide mixture was acidified and extracted from the gel as previously described^48,49^. Peptides were fractionated offline by high pH reverse-phase chromatography, loaded onto an EASY-Spray column (Thermo Fisher Scientific ES800: 15 cm × 75 μm ID, PepMap C18, 3 μm) via an EASY-nLC 1000, and gradient-eluted for online ESI-MS and MS/MS analyses with a Q Exactive Plus mass spectrometer (Thermo Fisher Scientific). MS/MS analyses of the top 8 precursors in each full scan used the following parameters: resolution: 17,500 (at 200 Th); AGC target: 2 × 105; maximum injection time: 800ms; isolation width: 1.4 m/z; normalized collision energy: 24%; charge: 3–7; intensity threshold: 2.5 × 103; peptide match: off; dynamic exclusion tolerance: 1,500 mmu. Cross-linked peptides were identified from mass spectra by pLink^50^. All spectra reported here were manually verified as previously^48^ and all cross-links are listed in Supplementary data table 1. Cross-links were visualized using xiNET^51^.

### Data availability

The cryo-EM density maps for the yeast SSU processome have been deposited in the EM Data Bank with accession codes EMD-AAAA, EMD-BBBB, EMD-CCCC, EMD-DDDD and EMD-EEEE. Corresponding atomic coordinates have been deposited in the Protein Data Bank under accession code XXXX.

## Supplementary Information

Supplementary information includes 36 Figures, two tables, one data table, one data file and one video.

## Acknowledgements

We thank M. Ebrahim and J. Sotiris for outstanding support with data collection at the Evelyn Gruss Lipper Cryo-EM resource center at The Rockefeller University. We further thank T. Walz, G. Alushin and Y. Shi for helpful discussions. J.B. is supported by an EMBO long-term fellowship (ALTF 51-2014) and a Swiss National Science Foundation fellowship (155515), M.C.-M. is supported by a postgraduate scholarship from the Natural Sciences and Engineering Research Council of Canada (NSERC). S.K. is supported by the Robertson Foundation, the Alfred P. Sloan Foundation, the Irma T. Hirschl Trust, the Alexandrine and Alexander L. Sinsheimer Fund, the Human Frontier Science Program Career Development Award, and an NIH New Innovator Award (1DP2GM123459). B.T.C. is supported by National Institute of Health Grant Nos. P41GM103314 BTC and P41GM109824 BTC.

## Author contributions

S.K. established purification conditions and M.C.-M. and J.B. acquired cryo-EM data. K.M. performed mass spectrometry experiments and analyzed the resulting data with B.C.. J.B., M.C.-M, M.H. and S.K. determined the cryo-EM structure of the yeast SSU processome, built the atomic model, interpreted the results and wrote the manuscript. All authors edited the manuscript.

## Author information

The authors declare no competing financial interests. Correspondence and requests for materials should be addressed to S.K.

